# Low-dimensional latent spaces identify the functional structure of individual behavioral phenotypes

**DOI:** 10.64898/2026.03.29.715160

**Authors:** Hiroshi Higashi

## Abstract

Extracting stable individual traits from behavior observed across diverse contexts is a central challenge in behavioral modeling. We propose a framework for inferring domain-invariant individual latent representations by jointly encoding behaviors across multiple domains. Using large-scale telemetry data from professional Counter-Strike 2 gameplay, we demonstrate that these representations are stable across distinct environments and roles, improving behavior prediction in novel domains. Our analysis reveals that complex idiosyncratic movement policies can be effectively compressed into low-dimensional embeddings, with as few as two dimensions capturing the majority of individual strategic variation. Crucially, the learned latent space forms a structured metric space where Euclidean distances predict the degradation of transfer performance. Furthermore, we show that the latent axes align with interpretable behavioral phenotypes, such as risk-taking and social cohesion. These findings suggest that multi-domain integration is a robust method for uncovering the functional structure of latent individuality in complex decision-making tasks, bridging the gap between high-dimensional telemetry data and meaningful psychological constructs.

## 1 Introduction

Understanding the latent factors that drive individual differences in behavior is a fundamental goal in cognitive science and artificial intelligence [1, 2]. While humans adapt their actions to specific environmental constraints and social roles, they often retain consistent behavioral “signatures” across diverse contexts. However, isolating these intrinsic traits from context-dependent adaptations remains a significant challenge. In most computational models, individual differences are often entangled with the specific structure of the environment or the requirements of a role, making it difficult to generalize behavior across unseen conditions. Effective behavioral modeling requires capturing individual variability through empirical priors that can be integrated into decision-making frameworks [3].

Traditionally, individual differences in behavioral style and strategic preference have been quantified through psychometric tools, such as self-report questionnaires, or highly controlled laboratory experiments [4, 5, 6, 7]. While these methods provide a foundation for identifying stable personality traits or cognitive profiles, they are often limited by subjective bias and low ecological validity when applied to the dynamic, high-dimensional nature of behavior in real-world or complex digital environments [8]. Recent computational approaches have begun to address these limitations by leveraging digital telemetry data—such as movement trajectories in video games or navigation patterns in virtual reality—to directly infer latent individual traits [9, 10, 11, 12, 13]. These data-driven methods allow for the discovery of behavioral “fingerprints” [10, 14] that are more representative of an individual’s actual decision-making style than what can be captured by traditional discrete scales.

Recent work on individuality transfer [14] has demonstrated that an encoder can extract an *individual latent representation* from behavior in one task condition and transfer it to predict behavior in another. However, extracting individuality from only a single *domain*—defined here as a specific task or role—may fail to distinguish between a person’s core tendencies and their reactions to that specific context. The paradox of statistical learning suggests that while mechanisms may be domain-general, they are often subject to modality-specific constraints [15]. To truly capture individuality, a model must identify patterns that persist across diverse domains, effectively disentangling intrinsic style from contextual demands.

In this study, we propose a multi-domain latent representation framework to address this limitation. Our main hypothesis is that jointly training a model to encode and decode behavioral evidence across multiple, distinct domains produces a more stable and “pure” representation of individuality than single-domain approaches. In our framework, a shared encoder integrates behavioral data from multiple source domains to infer a domain-robust latent code for each individual. This code is then used by domain-specific decoders (HyperNetworks [16]) to condition a behavior model on a target domain. By forcing a single latent code to serve as the basis for behavior prediction across radically different environments and strategic roles, the model is compelled to isolate the individual’s core behavioral signature from domain-specific adaptations. This approach builds on the principles of domain generalization [17] and meta-reinforcement learning [18], aiming to learn representations that remain invariant across diverse task conditions.

We employ Counter-Strike 2 (CS2) as a high-dimensional testbed for human decision-making [19]. CS2 offers a rich environment for testing the stability of extracted traits due to its diverse map geometries, distinct strategic roles, and the availability of large-scale competitive replay data. In this context, a domain is defined as a combination of a map’s spatial layout and the player’s side. By building individual behavior models that predict player trajectories, we demonstrate that a model conditioned on a multi-domain latent representation accurately reproduces behavior in unseen player–map–side combinations, effectively transferring learned individuality across radically different spatial and strategic contexts. Furthermore, we characterize the informational requirements and functional logic of these representations, showing that idiosyncratic behavioral policies can be compressed into a low-dimensional metric space that aligns with interpretable strategic phenotypes.

## 2 Results

### 2.1 Dataset and task setup

We assembled a dataset of round-level player trajectories from professional Counter-Strike 2 (CS2) matches on three common maps: *Dust II, Mirage*, and *Inferno*. Professional gameplay recordings (demos) were sourced from the HLTV database^1^ for the period between October and December 2025. The resulting dataset encompasses 268 matches and 5,756 rounds of high-level gameplay. A behavioral domain, denoted as *d* = (*m, s*), was defined as a unique combination of map geometry *m* ∈ ℳ and strategic role *s* ∈ {*T, CT*}. We identify each domain using a three-letter abbreviation, where the first two letters denote the map and the last letter denotes the side (e.g., DUT for Dust II × Terrorist and MIC for Mirage × Counter-Terrorist).

To ensure sufficient behavioral evidence for each player, we included only those who participated in multiple matches for each map–side combination. After filtering, we identified a cohort of 91 unique players (the “common players”) who met these criteria across all three maps. For each round, we extracted time-ordered 2D coordinates (*X, Y*) sampled at 1 Hz, along with the center of positions of their allies (Ally_*X*_, Ally_*Y*_).

For each player *i* and domain *d*, we randomly partitioned their matches into source 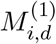 and target 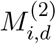 sets. This setup ensured that the behavioral data for the source and target domains were completely disjoint, allowing us to evaluate the stability of extracted traits when transferring from one map–side pair to another (e.g., from DUC to MIT).

### 2.2 A joint encoding–decoding framework for multi-domain individuality extraction

Our framework consists of five core components designed to isolate individual traits through joint optimization across diverse environmental and strategic contexts:

1. **Encoders**: Domain-specific encoders that process historical trajectories (60 s) from multiple source domains to extract latent representations. The representation is individually encoded for each source domain.
2. **Combiner**: A network component that integrates representations encoded from multiple source domains (e.g., via averaging) to form a domain-robust integrated representation. The combiner applies averaging to the incoming latent representations from the source domain’s encoders to output an integrated representation even if data from certain source domains is missing.
3. **Individual Latent Code (z)**: A *d*-dimensional vector acting as a stable, domain-invariant signature for the player. Our analysis primarily focused on a model with a 2-dimensional latent space (*d* = 2), which was selected based on preliminary hyperparameter tuning to balance representational capacity and generalization (Figure 5).
4. **Decoders (HyperNetworks)**: A set of domain-specific networks that map the shared latent code **z**_*i*_ to the parameter space of a trajectory predictor for each environment (map) and role (side).
5. **Trajectory Predictor**: A sequential model that predicts movement on a target domain, conditioned on the player’s extracted latent trait via the parameters generated by the corresponding decoder.

Figure 1 illustrates the flow of information through these components.

**Figure 1:**
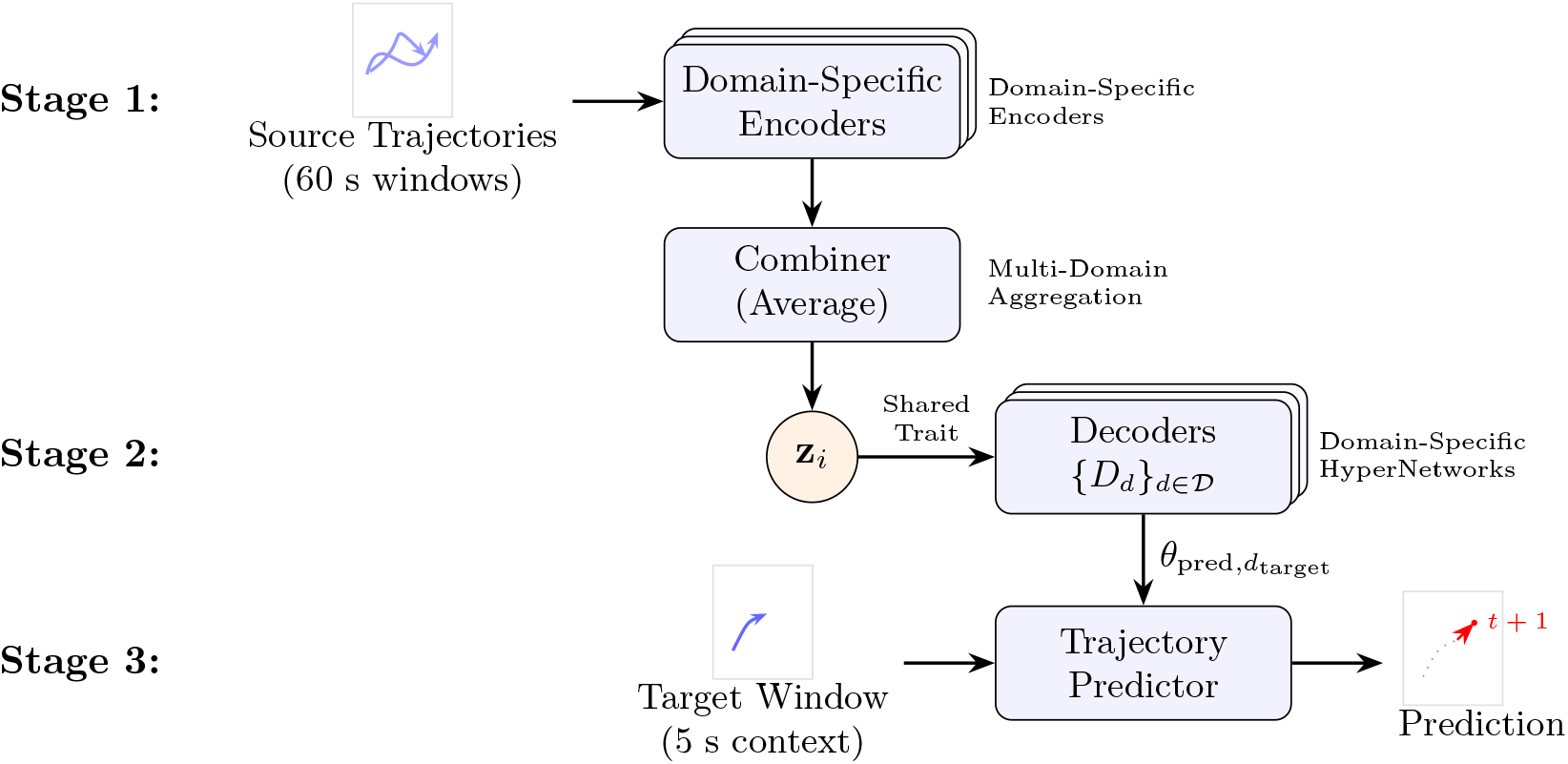
Multi-domain latent representation framework. Stage 1: Encoders process long-term source trajectories (60 s) from multiple domains, which are integrated by a Combiner. Stage 2: The domain-invariant latent code **z**_*i*_ is shared across domain-specific decoders. Stage 3: The decoder for the target domain generates parameters *θ*_pred_ for the trajectory predictor, which predicts the next-step position given a short-term (5 s) context.

By jointly training these components across all available map–side combinations, the model is forced to identify intrinsic strategic and movement styles that remain consistent regardless of the specific domain.

### 2.3 Multi-domain embeddings accurately specialize to individual movement signatures within a single domain

Prior to assessing cross-domain stability, we evaluated the capacity of the proposed framework to capture and utilize player-specific behavioral signatures within a consistent environment (i.e., a within-domain scenario). In this setting, the latent code **z** for each player was extracted from their historical trajectories in the same domain as the target domain, using data from an independent matchset (Matches *M* ^(1)^ for encoding **z**_*i*_ and *M* ^(2)^ for evaluation). Predictive performance was quantified using the root mean square error (RMSE) in raw coordinate units.

Our results demonstrate that the proposed multi-domain model effectively specializes its prediction logic to individual players when provided with historical data from the same environment. We found that both the choice of model and the specific domain significantly influenced predictive accuracy (two-way repeated measures analysis of variance (RM ANOVA), Model: *F*_3,36_ = 100.1, *p* < 0.001; Domain: *F*_5,60_ = 3.2, *p* = 0.012), with no significant interaction between the two (*F*_15,180_ = 1.4, *p* = 0.16). Post-hoc pairwise comparisons showed that the proposed model outperformed the population model—which represents the average behavior of the cohort—across all map domains (adjusted *p* < 0.001, Figure 2a). This improvement highlights the insufficiency of population-level models in capturing the nuanced movement styles of individual players, even within a single domain. Notably, the proposed model also achieved higher predictive accuracy than the Self baseline (*p* < 0.001), which was explicitly trained on each player’s specific data. While the Self model might theoretically serve as an upper bound through exhaustive optimization, these results suggest that individual-specific optimization is often constrained by data scarcity for single players—a limitation effectively mitigated by our multi-domain framework.

**Figure 2:**
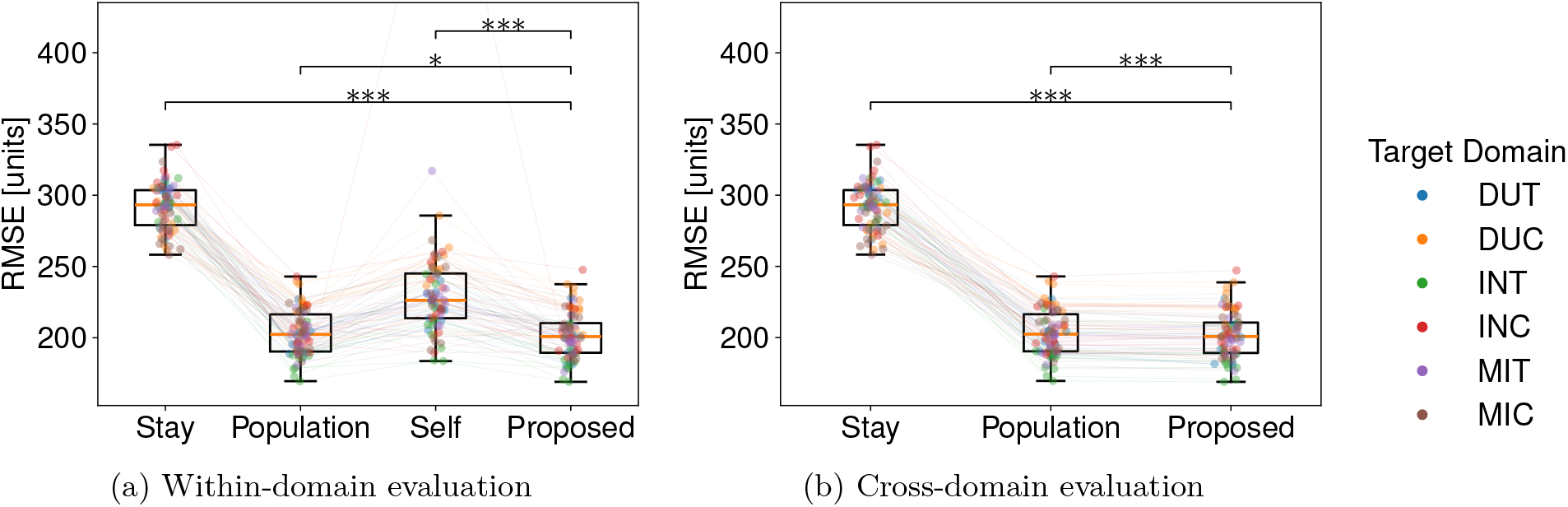
Comparison of trajectory prediction performance in the within-domain and cross-domain scenarios. The boxplots illustrate the distribution of RMSE for the compared models, where the central line indicates the median and the whiskers represent the non-outlier range. Individual data points (jittered dots) correspond to prediction error for each player, color-coded by the target domain. Asterisks denote statistical significance for comparisons between the proposed model and each baseline, determined by post-hoc *t*-tests with Bonferroni correction (^∗^*p* < 0.05, ^∗∗^*p* < 0.01, ^∗∗∗^*p* < 0.001). Outliers are not shown for clarity; vertical-axis scaling is based on the non-outlier data range (whiskers). (a) Within-domain scenario. The latent representation **z** for the proposed model was derived from historical data in the same domain as the target, allowing for direct specialization to individual movement styles. (b) Cross-domain scenario. The latent representation **z** for the proposed model was derived exclusively from historical data in domains other than the target domain, testing its ability to generalize player-specific movement patterns to unseen domains.

### 2.4 Extracted latent traits enable zero-shot individuality transfer across diverse environments

To investigate whether the extracted latent representations capture domain-invariant player signatures, we conducted a “cross-domain” evaluation. In this more challenging setting, we assessed the model’s ability to predict player trajectories in a completely novel domain. For a given target domain, the latent code **z** was computed by averaging the encoder outputs derived from the player’s historical data on all other available domains, excluding any data from the target domain itself.

Our cross-domain analysis revealed that the proposed framework successfully achieved zero-shot transfer by extracting domain-invariant behavioral features, consistent with the emerging use of foundation models in human behavior modeling [10]. In the cross-domain transfer task, model performance varied significantly across architectures and domains (two-way RM ANOVA, Model: *F*_2,24_ = 1377.1, *p* < 0.001; Domain: *F*_5,60_ = 3.9, *p* < 0.001), with a significant interaction indicating that some domains were inherently more challenging for transfer than others (*F*_10,120_ = 10.1, *p* < 0.001). Post-hoc comparisons demonstrated that the proposed model outperformed the population baseline across all novel domains (adjusted *p* < 0.001, Figure 2b), even in the complete absence of domain-specific historical data. These results suggest that the encoder successfully disentangled idiosyncratic movement styles from domain-specific constraints, projecting them into a latent space that generalizes across disparate spatial layouts and strategic roles. The performance gain relative to the population model indicates that individual movement patterns constitute a consistent behavioral signature that can be leveraged to enhance predictive accuracy in novel environments. Consequently, the latent embedding serves not merely as a record of historical locations but as a functional representation of an individual’s underlying behavioral policy.

Qualitative examples of trajectory predictions for representative players in unseen target domains are shown in Figure 3. While the proposed model significantly outperformed the population model on average, it did not consistently yield superior predictions in every instance.

**Figure 3:**
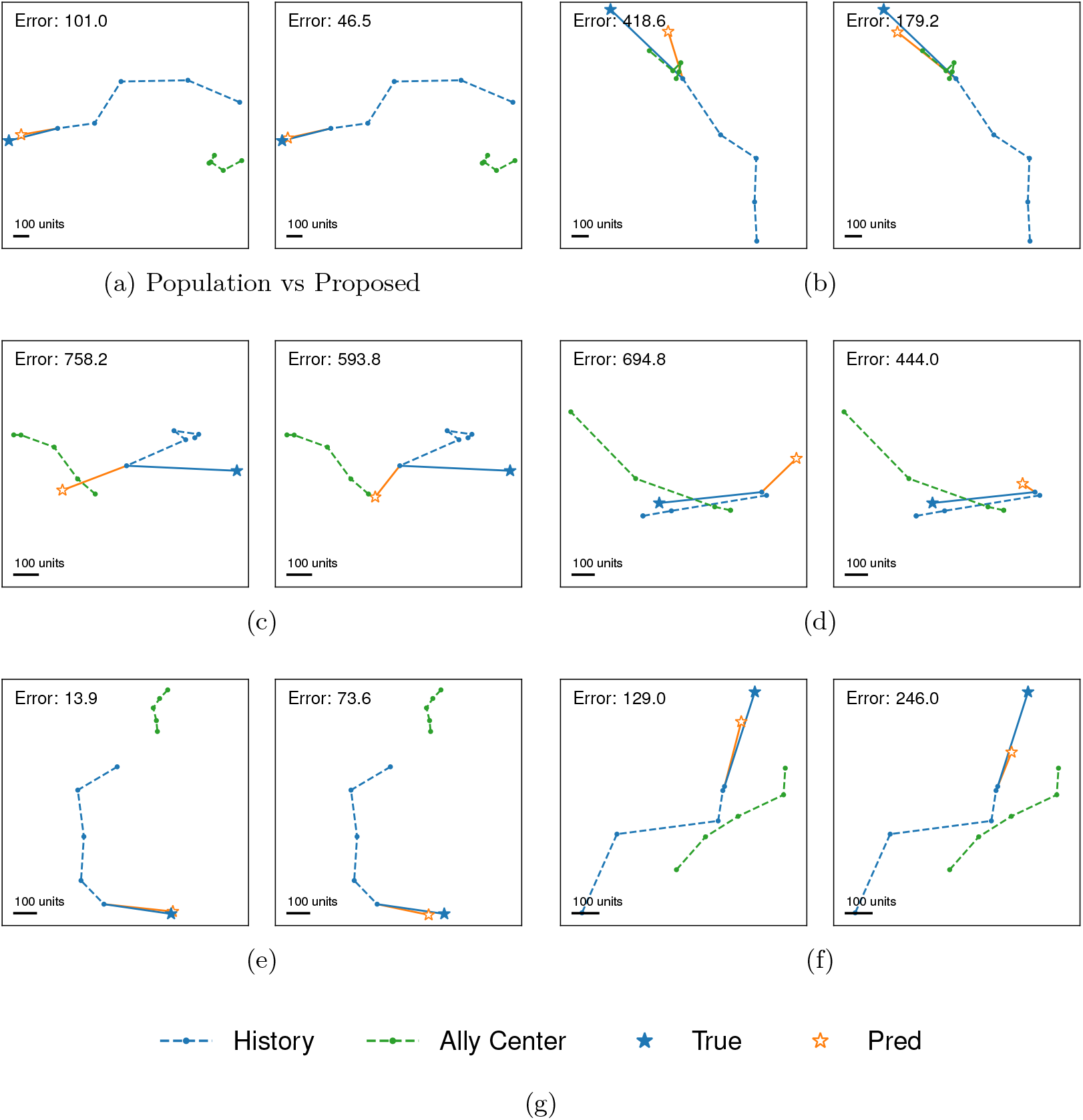
Qualitative comparison of trajectory predictions between the Population Model (left) and the Proposed Model (right) for representative players in unseen target domains. The 5-second movement history (dashed blue line) and the center of mass of allies (dashed green line) provide context for the prediction. (a, b) Examples where the proposed model’s prediction is more accurate than the population model. The population model tends to predict a position closer to the global mean of the training data, whereas the proposed model’s prediction (orange star) is more closely aligned with the ground truth (blue star). (c, d) Instances where both models deviate from the ground truth, but the proposed model provides a closer approximation. (e, f) Examples where the population model’s prediction aligns more closely with the actual behavior than the proposed model.

### 2.5 Aggregation of multi-source behavioral evidence enhances representation robustness

To assess how the availability of multi-source information influences the accuracy of zero-shot transfer, we performed a robustness analysis by varying the number of source domains (*k*) used to compute the latent representation **z**. For each target domain, we evaluated all possible combinations of *k* source domains (*k* ∈ {1, 2, 3, 4, 5}), selected from the historical data available in other environments.

Predictive performance was measured as relative RMSE, defined as the difference between the error at a given *k* and the error achieved with all available sources (*k* = 5). This normalization allowed us to isolate the marginal gain provided by each additional source environment. To statistically evaluate the effect of information diversity, we conducted a one-way RM ANOVA with Source Count (*k*) as the within-subject factor. Post-hoc comparisons between consecutive *k* values were performed using paired *t*-tests to identify the point at which performance improvements diminished.

Our analysis revealed that the diversity of source environments is a critical factor for successful individual adaptation in novel contexts. The diversity of source environments influenced predictive accuracy (one-way RM ANOVA, *F*_4,20_ = 12.1, *p* < 0.001). As illustrated in Figure 4, the prediction error decreased monotonically as more source domains were integrated into the latent embedding.

**Figure 4:**
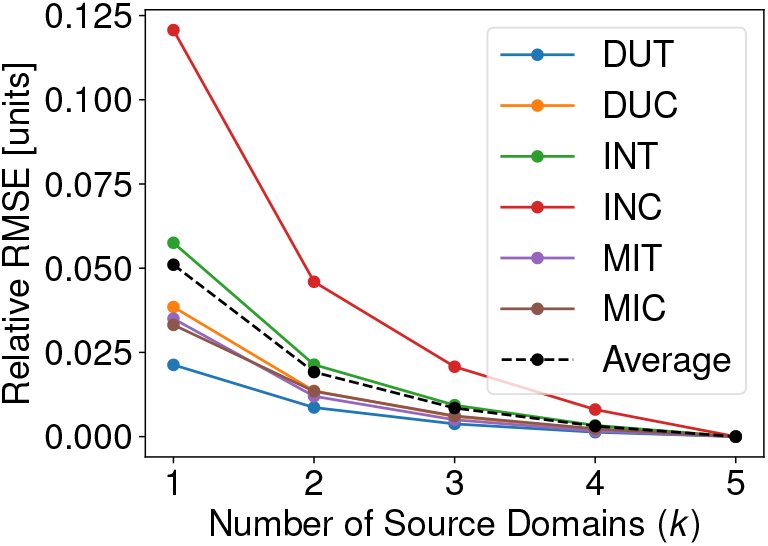
Robustness of zero-shot transfer to the number of source domains. The plot shows the relative RMSE as a function of the number of source environments (*k*) used to derive the latent vector **z.** For each target domain, the error at *k* = 5 is defined as the baseline (zero). Individual lines represent target maps, and the thick dashed line indicates the global average. Accuracy improves significantly as the diversity of source domains increases (*p* < 0.001, RM ANOVA), demonstrating the model’s ability to integrate complementary behavioral signatures across multiple domains.

**Figure 5:**
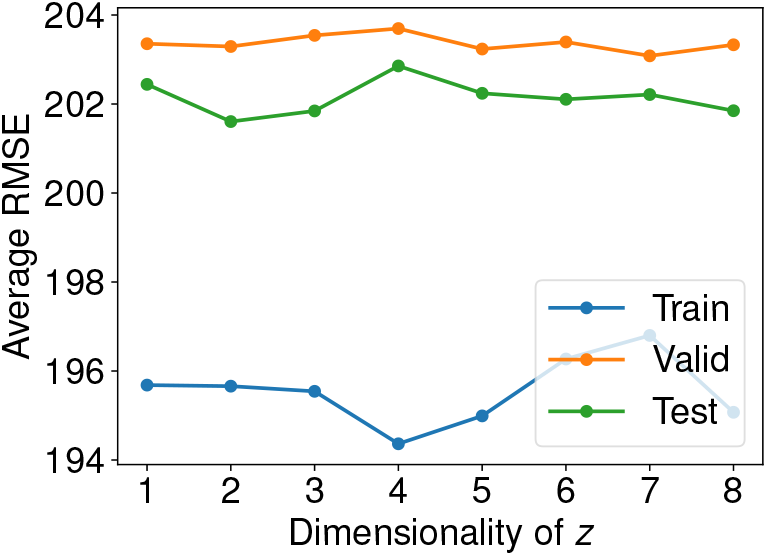
Influence of latent space dimensionality on predictive performance. The graph shows the average RMSE as a function of the dimensionality of the latent representation **z.** Results are presented for both the training, validation, and test datasets across all target domains. Error decreases as the dimensionality increases from 1 to 2, indicating that low-dimensional embeddings are sufficient to capture idiosyncratic behavioral signatures. Performance reaches a plateau beyond *d* = 2, suggesting an optimal balance between information compression and predictive accuracy.

Post-hoc analysis showed significant performance improvements when increasing from *k* = 1 to *k* = 2 (*p* = 0.017), with continued significant gains observed for each additional source up to *k* = 5 (all *p* < 0.05). This result demonstrates that diverse information is needed to form a more robust and accurate representation of the player’s behavioral policy. The consistent downward trend across all target domains suggests that the proposed framework is not dependent on specific domain pairs but instead benefits from a global increase in the “behavioral samples” provided by multiple environments.

### 2.6 Low-dimensional latent codes are sufficient to capture individual strategic variation

A key challenge in behavioral modeling is determining the informational complexity required to distinguish one individual’s strategic policy from another. To address this, we evaluated the model’s predictive performance while varying the dimensionality of the latent embedding **z** across *d* ∈ {1, …, 8}, while maintaining all other architectural parameters constant. The generalization capability of these representations was assessed using the average RMSE across all target domains for the training, validation, and test datasets (Matches *M* ^(2)^).

Our analysis revealed that while training error decreased with higher dimensionality, generalization performance on validation and test sets remained remarkably stable for *d* ≥ 2 (oneway RM ANOVA, Training: *F*_7,413_ = 51.9, *p* < 0.001; Validation: *F*_7,84_ = 0.6, *p* = 0.771; Test: *F*_7,84_ = 2.8, *p* = 0.010). As illustrated in Figure 5, a single-dimensional representation (*d* = 1) was insufficient to capture the complexity of movement strategies, resulting in higher error. However, performance reached a stable plateau from *d* = 2 onwards, suggesting that additional dimensions provide marginal gains in generalization accuracy relative to the increase in representational complexity.

To further characterize the functional structure of these representations, we visualized the distribution of the learned latent vectors (**z**) for *d* = 2 and *d* = 4 across different data splits (Figure 6). Crucially, the latent representations of individuals from the training, validation, and test sets occupied a consistent region of the latent space with high overlap. This consistency indicates that the encoder successfully learned a generalized mapping of behavioral styles that extends to novel individuals, rather than over-fitting to the specific players in the training set. Moreover, pairwise visualization of the latent space revealed that individuals are distributed along a continuous manifold without clear clusters. This topology suggests that player “style” in competitive environments is best described by a set of continuous behavioral gradients—such as varying degrees of risk-taking or social cohesion—rather than discrete strategic archetypes.

**Figure 6:**
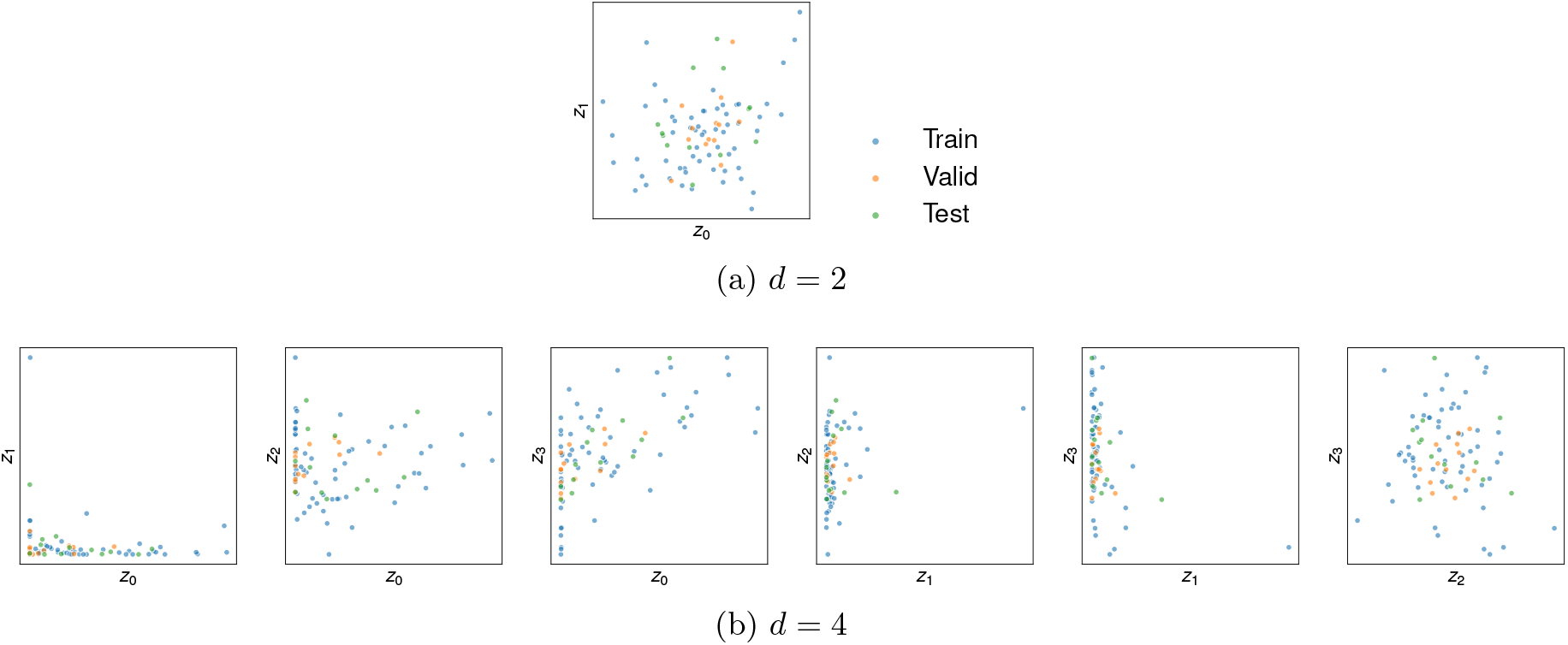
Representative pairwise distributions of latent vectors (**z**) for *d* = 2 (top) and *d* = 4 (bottom). Each dot represents the latent signatures of an individual player, color-coded by data split. The high degree of overlap between splits demonstrates the model’s ability to map novel individuals into a stable, pre-defined behavioral manifold. Axis ticks are omitted to emphasize the relative distribution and topology of the latent space.

These findings demonstrate that complex behavioral phenotypes can be effectively compressed into a highly condensed latent code without sacrificing generalization performance. Based on this evidence, we adopted a 2-dimensional latent space (*d* = 2) for all subsequent analyses as a parsimonious representation of individual strategic variation. This low-dimensional compression aligns with the principles of information theory and recent work on disentangled representations, where complex generative factors are captured within a compact metric space [20, 21].

### 2.7 The learned latent space forms a structured metric manifold of behavioral policies

To verify whether the learned latent space preserves the metric relationships between different behavioral phenotypes, we conducted a cross-player transfer analysis. We reasoned that if the latent representation **z** effectively captures idiosyncratic behavioral policies, prediction error for a target player *i* should increase as the surrogate latent vector **z**_*j*_ used for model conditioning deviates from the player’s true representation **z**_*i*_.

For each map domain, we extracted reference latent vectors for all players from Matchset *M* ^(1)^. For a representative subset of target players, we performed trajectory predictions using the latent vectors of all other players in the same domain. To isolate the effect of latent distance from individual differences in baseline predictability, we defined relative RMSE as the difference between the error obtained using a surrogate vector **z**_*j*_ and the baseline error using the player’s own vector **z**_*i*_ (where ∥**z**_*i*_ − **z**_*j*_∥ = 0).

We found that the topological structure of the learned latent variable spaces was highly informative of behavioral similarity, which is a key property of models using amortized inference to capture cognitive dynamics [22]. A linear mixed-effects model (LMM) analysis—treating latent distance as a fixed effect and target player nested within domain as a random effect—revealed a significant positive relationship between latent distance and the degradation in predictive performance (fixed effect *β* = 51.26, *p* < 0.001; Figure 7). This metric structure suggests that the encoder organizes player signatures in a way that reflects their underlying behavioral policies, potentially forming a manifold of strategic phenotypes [23].

**Figure 7:**
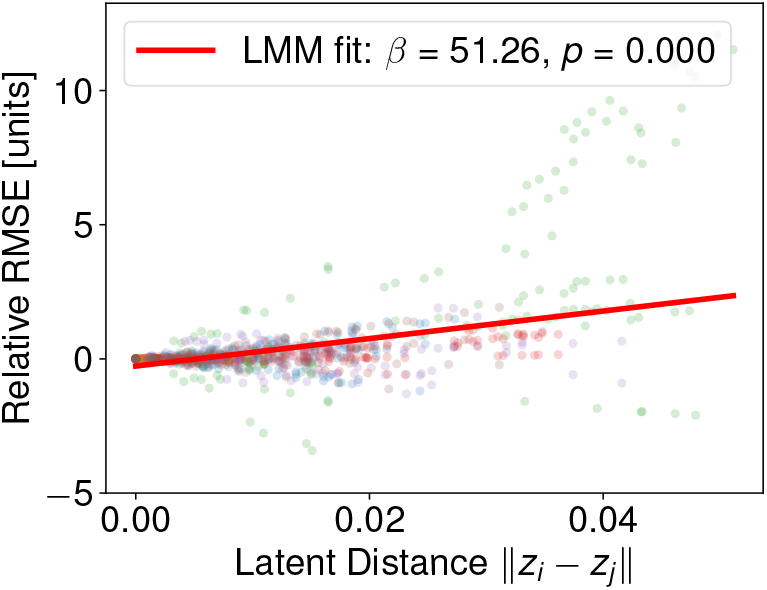
Metric structure of the latent space correlates with behavioral dissimilarity. The scatter plot illustrates the relationship between the Euclidean distance between two players in the latent space (∥**z**_*i*_ − **z**_*j*_∥) and the resulting change in predictive performance (relative RMSE). Relative RMSE was calculated by predicting the trajectory of a target player *i* using the latent vector **z**_*j*_ of a surrogate player *j*, normalized by the player’s own baseline error. Data points are color-coded by domain. The red line indicates the fixed-effect trend estimated by an LMM, accounting for individual and domain-specific variability. The positive slope (*β*) and significant *p*-value demonstrate that proximity in the latent space effectively represents phenotypic similarity in movement behavior.

As shown in the distance-accuracy gradient (Figure 7), surrogate latent vectors that were closer to the target player’s true representation yielded substantially lower prediction errors than more distant vectors. This relationship indicates that the encoder does not merely assign arbitrary identities to players. Instead, it projects idiosyncratic movement traits into a structured metric space where the HyperNetwork-based adaptation mechanism successfully leverages these low-dimensional behavioral signatures to modulate the predictor’s logic in a consistent and interpretable manner.

### 2.8 Latent axes align with interpretable strategic phenotypes like risk-taking and team cohesion

To elucidate the functional role of the learned latent space, we investigated the relationship between the low-dimensional embeddings **z** and observable behavioral phenotypes. We extracted eight key behavioral metrics for each player from the raw game logs, including average movement speed, visual scanning rate (yaw change), vertical mobility, positioning dispersion, survival duration per round, and average proximity to teammates. These metrics were aggregated across 532 independent matches to ensure a robust representation of each player’s characteristic style. The histograms for these metrics are shown in Figure 9.

We focused our interpretability analysis on a model with a two-dimensional latent space (**z** ∈ ℝ^2^) to identify the primary axes of behavioral variation. For each player identified in both the tracking datasets and the behavioral logs (*N* = 91), we calculated their mean latent vector across all map domains. The association between each latent dimension (*z*_0_, *z*_1_) and the behavioral metrics was quantified using Spearman’s rank correlation coefficient (*ρ*). All *p*-values were adjusted for multiple comparisons where appropriate, and a significance threshold of *α* = 0.05 was used.

The two-dimensional latent space effectively captured distinct facets of player behavior, providing an interpretable basis for the predictor’s adaptation logic. Our correlation analysis revealed that the latent dimensions were not merely abstract identities but were significantly aligned with specific behavioral traits (Figure 8).

**Figure 8:**
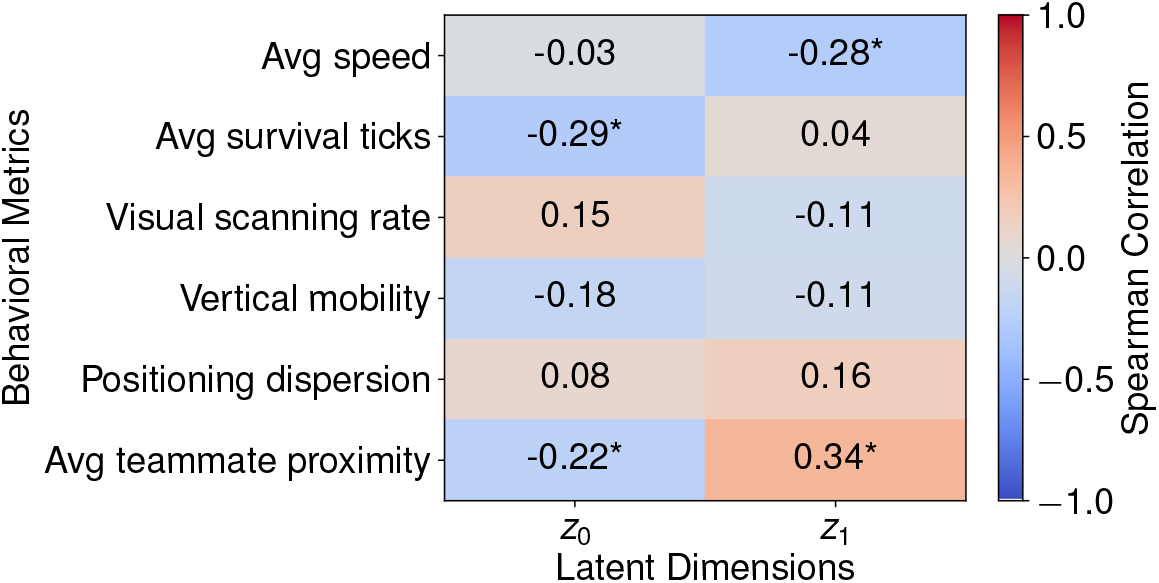
Correlation between latent dimensions and behavioral metrics. The heatmap displays Spearman’s rank correlation coefficients (*ρ*) between each dimension of the latent embedding (*z*_0_ and *z*_1_) and various behavioral metrics extracted from game logs. Asterisks indicate statistically significant correlation ^∗^*p* < 0.05). The color scale represents the strength and direction of the correlation, ranging from −1 (dark blue) to +1 (dark red). The results demonstrate that the learned latent space naturally encodes interpretable behavioral phenotypes, such as survival duration and team coordination tendencies.

**Figure 9:**
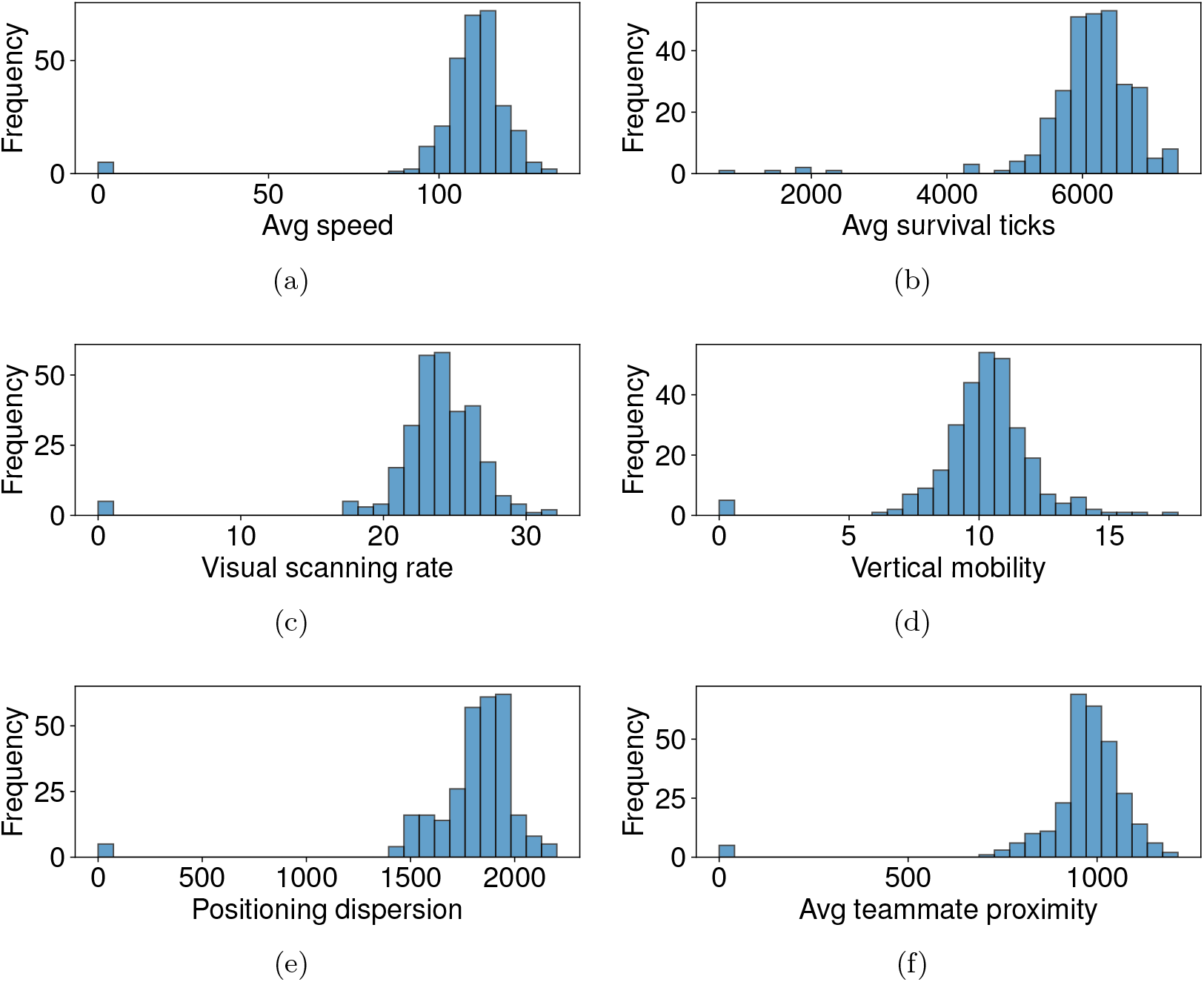
Distributions of behavioral metrics across players. Histograms for each of the eight behavioral metrics extracted from game logs, aggregated across all players and matches. These distributions provide context for the range and variability of behaviors captured by the latent space, illustrating the diversity of player strategies in the dataset.

Specifically, the first latent dimension (*z*_0_) was primarily associated with survival duration and team cohesion. We observed a significant negative correlation between *z*_0_ and average survival ticks (*ρ* = −0.29, *p* = 0.006), as well as average teammate proximity (*ρ* = −0.22, *p* = 0.035; Figure 10a, b). This suggests that *z*_0_ might represent a spectrum between high-risk, aggressive solo-play and more conservative, team-oriented strategies.

**Figure 10:**
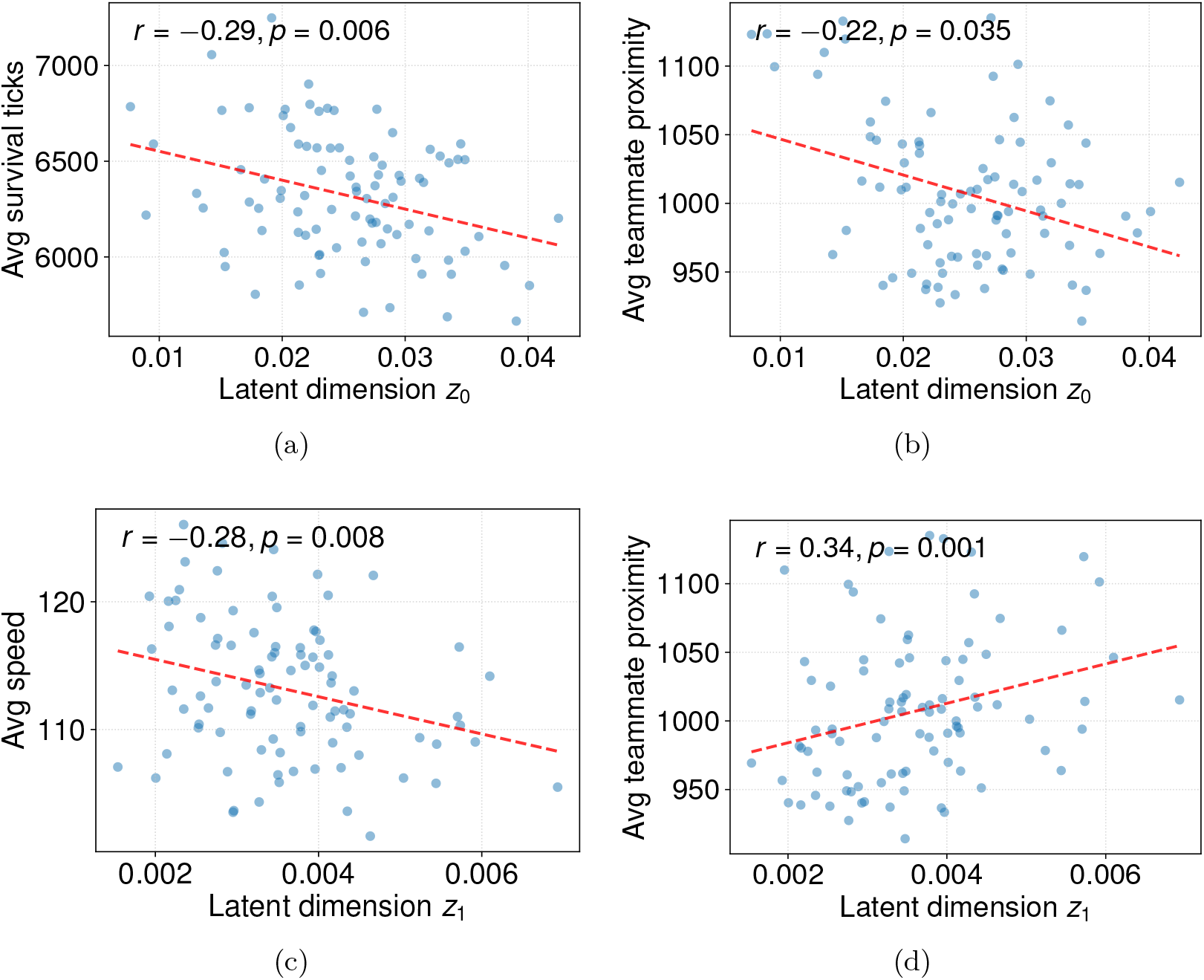
Detailed relationships for significant latent-behavior pairs. Scatter plots illustrate the linear relationships for pairs of latent dimensions and behavioral metrics that exhibited significant correlations (*p* < 0.05). Each point represents an individual player (*N* = 91). The dashed red lines indicate linear regression fits, with Spearman’s correlation coefficient (*r*) and *p*-value provided in each panel. These plots highlight how specific latent axes track fine-grained behavioral variations, such as the trade-off between movement speed and teammate proximity or the predictors of survival efficiency.

The second dimension (*z*_1_) demonstrated a strong relationship with mechanical movement intensity and cooperative positioning. *z*_1_ showed a significant negative correlation with average movement speed (*ρ* = −0.28, *p* = 0.008) and a significant positive correlation with teammate proximity (*ρ* = 0.34, *p* = 0.001; Figure 10c, d). These results indicate that *z*_1_ effectively disentangles fast-paced individualistic movement from slower, coordinated group movements.

The emergence of such interpretable axes within an end-to-end trained transfer learning framework suggests that the HyperNetwork-based architecture naturally organizes player representations according to functionally relevant behavioral phenotypes. This organization allows the model to modulate its internal logic based on fundamental strategic differences between players, thereby achieving superior adaptive performance.

## 3 Discussion

The key contribution of this work is a novel framework for extracting individual behavioral traits from multi-domain data, where domains are defined by both environmental (map) and strategic (side) contexts. While humans are famously adaptable to different roles and environments, we have shown that a core latent individuality can be isolated from these context-dependent actions. Our results demonstrate that this extraction is significantly more robust when evidence is aggregated across multiple, distinct contexts and roles, with predictive accuracy improving monotonically as the diversity of source environments increases.

These findings have implications for the study of individual differences in behavioral science. Rather than viewing individuality as a set of fixed parameters within a single task or role, our multi-domain approach allows us to define individuality by its consistency across diverse challenges. We found that the informational complexity of human movement signatures is relatively low, with low-dimensional embeddings (*d* = 2) sufficient to capture the nuances of individual strategic policies. Crucially, the learned latent space is not merely an abstract identification system but a structured metric space where distances reflect behavioral dissimilarity. The emergence of interpretable latent axes—correlating with fundamental traits such as risk-taking (survival duration) and social cohesion (teammate proximity)—suggests that our framework identifies stable “behavioral phenotypes” that bridge the gap between high-dimensional telemetry data and meaningful psychological constructs. These phenotypes provide a basis for adaptive behavior, integrating experience and instruction to refine individual strategies over time [24]. The discovery of such stable, domain-invariant signatures represents a step toward modeling the lifelong learning and knowledge transfer characteristic of biological systems, a fundamental challenge for contemporary artificial intelligence [25].

Although CS2’s behaviors (trajectories) are highly contextual, the player’s roles are mostly fixed because the game is team-based. Therefore, the extracted traits may partly reflect the strategic requirements of these roles, rather than the purely intrinsic nature of the individual players. Nevertheless, the fact that our framework successfully extracts consistent behavioral patterns in an environment with such strictly defined roles demonstrates its ability to uncover latent play tendencies directly from trajectory data.

Another consideration is that our current implementation focuses exclusively on trajectory prediction. While we demonstrate that individual strategic signatures are robust across environments and roles, these representations were optimized for spatial navigation. The underlying framework is inherently task-agnostic; in principle, the same individual latent codes could be used to condition models for other high-dimensional behavioral modalities, such as gaze patterns or tactical decision-making [10]. Validating the stability of these traits across such heterogeneous task types remains a critical objective for future research to further establish the domain-invariance of human behavioral phenotypes.

Looking forward, our framework provides a foundation for several lines of inquiry into the functional architecture of human strategic behavior. First, elucidating the alignment between data-driven latent traits and established psychometric profiles will provide a quantitative bridge between high-dimensional behavioral telemetry and formalized personality theory. Second, our multi-domain approach offers a promising avenue for investigating collective dynamics, specifically how idiosyncratic strategic phenotypes interact to determine team-level synergy and collective intelligence [26]. Finally, the ability to extract stable, domain-invariant signatures from behavioral evidence presents a robust tool for building high-fidelity personalized models. Beyond the scope of competitive gaming, such personalized modeling holds significant potential for broader societal impact, ranging from the development of individual-specific biomarkers for cognitive monitoring to the design of adaptive, user-centric systems that respond to the unique behavioral styles of their users [27, 28].

## 4 Materials and methods

### 4.1 Data preprocessing

Counter-Strike 2 demo files were parsed into tick-level logs using the community tool `demoinfocs-golang` [29]. Trajectories were resampled to 1 Hz to capture macro-level movement patterns. For each player, we extracted 2D coordinates (*X, Y*), role *s*, and the mean position of their four teammates (Ally_*X*_, Ally_*Y*_) as a social context feature. Coordinates were normalized using common statistics across all maps (mean *µ* = [0, 500, 0, 500], std *σ* = [1500, 1500, 1500, 1500]) to ensure a consistent coordinate frame. We focused on three major competitive maps: *Dust II, Mirage*, and *Inferno*.

A domain *d* is defined as a specific (map, side) pair. To ensure the stability of the extracted traits, we selected players who participated in at least two matches for each of the three maps. For each player–map combination, matches were randomly partitioned into two sets: *M* ^(1)^ used for source domain evidence and *M* ^(2)^ used for target domain evaluation. The players were then split into training (70%), validation (15%), and testing (15%) sets to evaluate the framework’s ability to generalize to unseen individuals, following common practices in domain generalization [17, 30].

### 4.2 Model details

The proposed framework employs an encoder–decoder architecture where the decoders are implemented as HyperNetworks [16]. Table 1 summarizes the architecture and hyperparameters of each component.

**Table 1:** Architectural details and hyperparameters of the multi-domain latent representation framework.

| Component | Architecture | Parameters/Dimensions |
| --- | --- | --- |
| **Encoder** | GRU [31] + 4-layer FC | Input: 60 s window ($60 \times 4$)<br>GRU Hidden: 64<br>Output ($\mathbf{z}_i$): $d$-dimensional |
| **Combiner** | Averaging layer | Input: $N \times d$-dimensional representations<br>Output: $d$-dimensional integrated code |
| **Decoder**<br>(HyperNetwork) | 4-layer FC | Input: $d$-dimensional latent code<br>Output: $\theta_{\text{pred}}$ (26,050 params) |
| **Trajectory Predictor** | GRU + 4-layer FC | Input: 5 s window ($5 \times 4$)<br>GRU Hidden: 64<br>Output: 2D position |

The Encoder processes a 60-second trajectory window (1 Hz × 60 s = 60 steps) from source domains. It consists of a single-layer GRU (64 hidden units) followed by a four-layer fully connected network that compresses the temporal information into a *d*-dimensional individual latent code **z**_*i*_. A Combiner merges latent vectors from multiple source domains using a simple averaging operation to form a domain-robust representation.

To handle diverse environmental and strategic contexts, we employ a set of Decoders, one for each target domain. Each Decoder is a HyperNetwork consisting of a four-layer fully connected network that takes the shared latent code **z**_*i*_ as input and generates the entire parameter set *θ*_pred,*d*_ for the trajectory predictor specific to domain *d*. By jointly training these domain-specific decoders with a shared encoder, the model is forced to capture intrinsic behavioral tendencies in **z**_*i*_ that are informative across all domains. The Trajectory Predictor is a sequential model (single-layer GRU with 64 hidden units and a four-layer fully connected network) that predicts the player’s position at the next second (*t* + 1), conditioned on the previous 5-second trajectory window and the generated parameters *θ*_pred,*d*_.

Training was performed for 20,000 epochs using the Adam optimizer [32] with a constant learning rate of 1 × 10^−5^. The models were trained using a player-wise full-batching strategy [33], where all available trajectory sequences for a given player in the target domain were processed simultaneously in a single forward pass. The objective function was the sum of mean squared errors (MSE) between the predicted and actual normalized coordinates across all target domains during the joint training phase. When calculating the loss for a target domain, we excluded the latent representation from the matching source domain to compute the combined code **z**_*i*_; for example, data from a specific role in a given map was not used to predict behavior in independent matches of the same player– environment–role combination.

To ensure that the framework is robust to the absence of data from certain environments, we implemented a stochastic source dropout mechanism during training. For each training batch, a random number of available source domains (excluding the target domain) were omitted before computing the integrated latent code **z**_*i*_. This procedure forces the combiner and decoders to rely on a subset of behavioral evidence, encouraging the model to extract more generalized, domain-invariant traits rather than over-fitting to specific domain combinations. Consequently, the model maintains high predictive performance even when only limited historical data is available for a new player.

### 4.3 Statistical analysis

To evaluate the predictive accuracy across different models and domains, we primarily used the root mean square error (RMSE) in raw coordinate units. Statistical significance was determined using repeated measures ANOVA (RM ANOVA) with Model and Domain as within-subject factors for the within-domain and cross-domain comparisons. Post-hoc pairwise comparisons were performed using paired *t*-tests with Bonferroni correction for multiple comparisons. For the dimensionality analysis, a one-way RM ANOVA was conducted with latent dimensionality *d* as the within-subject factor. The relationship between latent distance and predictive performance was assessed using a linear mixed-effects model (LMM), with latent distance as a fixed effect and target player nested within domain as a random effect. Associations between latent dimensions and behavioral metrics were quantified using Spearman’s rank correlation coefficient (*ρ*), with *p*-values adjusted for multiple comparisons where appropriate. All statistical tests were performed with a significance threshold of *α* = 0.05.

### 4.4 Baselines and ablations

The performance of the proposed multi-domain framework was benchmarked against the following baselines:

1. **Stay model**: A naive baseline that predicts the next position (*t* + 1) to be identical to the current position (*t*). This model represents a zero-velocity assumption and serves as a fundamental reference for predictive accuracy.
2. **Population model**: A domain-specific trajectory predictor trained on the pooled behavioral data of all players in the training set for a given map–side combination. This model captures the average strategic movement patterns within a specific domain but lacks any mechanism for individual adaptation.
3. **Self model**: A personalized baseline where a separate trajectory predictor was trained specifically for each individual player using their own data from Matchset *M* ^(1)^. This serves as a theoretical upper bound for individual specialization; however, its performance is often limited by the data scarcity inherent in single-player datasets.
4. **Single-domain trait extraction (Ablation)**: An ablation of the proposed framework where the individual latent code **z**_*i*_ is derived from behavioral evidence in only a single source domain. This variant was used to quantify the informational benefit of aggregating trait evidence across diverse environments and strategic roles.

## 5 Data and code availability

Processed trajectory features, training scripts, and evaluation code will be released at publication. Raw demos are available from official match archives subject to platform terms.

## 6 Acknowledgements

This work was supported in part by the Japan Society for the Promotion of Science (JSPS) KAKENHI, grant numbers 22H05163 and 24K15047, and Japan Science and Technology Agency (JST) Advanced International Collaborative Research Program (AdCORP), grant number JPMJKB2307.

## 7 Author contributions

Conceptualization, Methodology, Software, Validation, Writing – original draft, Writing – review and editing: Hiroshi Higashi.

## A Supplementary Figures

## Footnotes

1 https://www.hltv.org/

## References

[1] Kriegeskorte N, Douglas PK. Cognitive computational neuroscience. Nat Neurosci. 2018;21(9):1148–60.

[2] Kass RE, Amari SI, Arai K, Brown EN, Diekman CO, Diesmann M, et al. Computational Neuroscience: Mathematical and Statistical Perspectives. Annu Rev Stat Appl. 2018 Mar;5(1):183–214.

[3] Gershman SJ. Empirical priors for reinforcement learning models. J Math Psychol. 2016;71:1–6.

[4] Boogert NJ, Madden JR, Morand-Ferron J, Thornton A. Measuring and understanding individual differences in cognition. Philos Trans R Soc Lond B Biol Sci. 2018 Sep;373(1756):20170280.

[5] Costa PT Jr, McCrae RR. Four ways five factors are basic. Pers Individ Dif. 1992 Jun;13(6):653–65.

[6] Cattell RB. The Scientific Analysis of Personality. Rothe JP, editor. London, England: Routledge; 2017.

[7] Eysenck HJ. The Biological Basis of Personality. Charles C. Thomas Publisher; 1977.

[8] Chaytor N, Schmitter-Edgecombe M. The ecological validity of neuropsychological tests: A review of the literature on everyday cognitive skills. Neuropsychol Rev. 2003 Dec;13(4):181–97.

[9] Zook A, Fruchter E, Riedl MO. Automatic playtesting for game parameter tuning via active learning. arXiv [csAI]. 2019 Aug.

[10] Xie Y, Li Z, Wang X, Pan Y, Liu Q, Cui X, et al. Be.FM: Open Foundation Models for Human Behavior. arXiv preprint. 2025 May.

[11] Lu Z, Golomb J. Human EEG and artificial neural networks reveal disentangled representations and processing timelines of object real-world size and depth in natural images. Elife. 2025 Dec;13(RP98117).

[12] Drachen A, Seif El-Nasr M, Canossa A. Game Analytics – The Basics. In: Game Analytics. London: Springer London; 2013. p. 13–40.

[13] Drachen A, Sifa R, Bauckhage C, Thurau C. Guns, swords and data: Clustering of player behavior in computer games in the wild. In: 2012 IEEE Conference on Computational Intelligence and Games (CIG). IEEE; 2012. p. 163–70.

[14] Higashi H. Predicting human decision-making across task conditions via individuality transfer. Elife. 2026 Jan;14(RP107163).

[15] Frost R, Armstrong BC, Siegelman N, Christiansen MH. Domain generality versus modality specificity: The paradox of statistical learning. Trends Cogn Sci. 2015;19(3):117–25.

[16] Ha D, Dai A, Le QV. Hypernetworks. arXiv preprint. 2016 Sep;(arXiv:1609.09106).

[17] Dissanayake T, Fernando T, Denman S, Ghaemmaghami H, Sridharan S, Fookes C. Domain Generalization in Biosignal Classification. IEEE Trans Biomed Eng. 2020;68(6):1978–89.

[18] Wang JX, Kurth-Nelson Z, Tirumala D, Soyer H, Leibo JZ, Munos R, et al. Learning to reinforcement learning. arXiv preprint. 2017 Jan;(arXiv:1611.05763).

[19] Yannakakis GN, Togelius J. Artificial Intelligence and Games. Cham: Springer Nature Switzerland; 2025.

[20] Shannon CE. A mathematical theory of communication. Bell Syst Tech J. 1948;27(3):379–423.

[21] Burgess CP, Higgins I, Pal A, Matthey L, Watters N, Desjardins G, et al. Understanding disentangling in beta-VAE. arXiv preprint. 2018 Apr;(arXiv:1804.03599).

[22] Pan TF, Li JJ, Thompson B, Collins A. Latent Variable Sequence Identification for Cognitive Models with Neural Network Estimators. arXiv preprint. 2024 Jun.

[23] Shepard RN. Multidimensional scaling, tree-fitting, and clustering. Science. 1980 Oct;210(4468):390–8.

[24] Schiffer AM, Siletti K, Waszak F, Yeung N. Adaptive behaviour and feedback processing integrate experience and instruction in reinforcement learning. Neuroimage. 2017;146:626–41.

[25] Parisi GI, Kemker R, Part JL, Kanan C, Wermter S. Continual lifelong learning with neural networks: A review. Neural Netw. 2019 May;113:54–71.

[26] Woolley AW, Chabris CF, Pentland A, Hashmi N, Malone TW. Evidence for a collective intelligence factor in the performance of human groups. Science. 2010 Oct;330(6004):686–8.

[27] Yamada T, Hashimoto RI, Yahata N, Ichikawa N, Yoshihara Y, Okamoto Y, et al. Resting-state functional connectivity-based biomarkers and functional mri-based neurofeedback for psychiatric disorders: A challenge for developing theranostic biomarkers. Int J Neuropsychopharmacol. 2017;20(10):769–81.

[28] Lotte F, Bougrain L, Cichocki A, Clerc M, Congedo M, Rakotomamonjy A, et al. A review of classification algorithms for EEG-based brain-computer interfaces: A 10 year update. J Neural Eng. 2018;15(3):aab2f2.

[29] demoinfocs-golang: A Counter-Strike 2 Demo Parser for Go (demoinfo);. Accessed: 2026-3-25. https://github.com/markus-wa/demoinfocs-golang.

[30] Dockès J, Varoquaux G, Poline JB. Preventing dataset shift from breaking machine-learning biomarkers. Gigascience. 2021 Sep;10(9).

[31] Cho K, van Merrienboer B, Gulcehre C, Bahdanau D, Bougares F, Schwenk H, et al. Learning phrase representations using RNN encoder–decoder for statistical machine translation. In: Moschitti A, Pang B, Daelemans W, editors. Proceedings of the 2014 Conference on Empirical Methods in Natural Language Processing (EMNLP). Stroudsburg, PA, USA: Association for Computational Linguistics; 2014. p. 1724–34.

[32] Kingma DP, Ba J. Adam: A method for stochastic optimization. arXiv [csLG]. 2014 Dec.

[33] Paszke A, Gross S, Massa F, Lerer A, Bradbury J, Chanan G, et al. PyTorch: An imperative style, high-performance deep learning library. arXiv [csLG]. 2019 Dec.

